# Species composition and risk of transmission of *Aedes*-borne arboviruses around some Yellow hotspot areas in Northern Ghana

**DOI:** 10.1101/2020.06.02.129460

**Authors:** Joannitta Joannides, Mawuli Dzodzomenyo, Faustus Azerigyik, Eudocia Esinam Agbosu, Deborah Pratt, Joseph Osei Nyarkoh, Rebecca Pwalia, Godwin Kwame Amlalo, Maxwell A. Appawu, Hayashi Takashi, Shiroh Iwanaga, Andrea Buchwald, Rosemary Rochford, Daniel Boakye, Kwadwo Koram, Kofi Bonney, Samuel Kweku Dadzie

**Affiliations:** Department of Parasitology, Noguchi Memorial Institute for Medical Research, University of Ghana, Ghana; Department of Environmental and Occupational Health, School of Public Health, University of Ghana, Ghana; Department of Virology, Noguchi Memorial Institute for Medical Research, University of Ghana, Ghana; Department of Epidemiology, Noguchi Memorial Institute for Medical Research, University of Ghana, Ghana; Department of Environmental and Occupational Health, School of Public Health, University of Colorado, Anschutz Medical Campus, Aurora, CO, United States; Department of Immunology and Microbiology, University of Colorado, Anschutz Medical Campus, Colorado, United States; Department of Molecular Virology, Tokyo Medical and Dental University, 1-5-45, Bunkyo-ku, Tokyo 113-8510, Japan; Department of Environmental Parasitology, Tokyo Medical and Dental University, 1-5-45, Bunkyo-ku, Tokyo 113-8510, Japan

## Abstract

*Aedes*-borne viral diseases mainly Yellow Fever (YF), Dengue (DEN), Zika (ZIK) and Chikungunya (CHK) have contributed to many deaths’ in the world especially in Africa. There have been major outbreaks of these diseases in West Africa. YF outbreaks have occurred in Ghana. Although Ghana has not recorded any outbreak of DEN, ZIK and CHK, the risk is high due to its proximity to West African countries where outbreaks have been recently been recorded. This study assessed the risk of transmission of Yellow fever (YFV), Dengue (DENV), Chikungunya (CHKV) and Zika (ZIKV) viruses in Larabanga and Mole Game Reserve areas in Northern Ghana. The immature and adult stages of *Aedes* mosquitoes were collected from Larabanga and Mole Game Reserve area. There was a significant (P>0.001) number of mosquitoes collected during the rainy season than the dry season. A total of 1,930 *Aedes* mosquitoes were collected during the rainy season and morphologically identified. Of these, 1,915 (99.22%) were *Aedes aegypti* and 15 (0.22%) were *Aedes vittatus*. During the dry season, 27 *Aedes aegypti* mosquitoes were collected. A total of 415 *Ae. aegypti* mosquitoes were molecularly identified to subspecies level of which *Aedes (Ae) aegypti aegypti* was the predominant subspecies. Both *Ae. aegypti aegypti and Ae aegypti formosus* exist in sympatry in the area. All *Aedes* pools (75) were negative for DENV, ZIKV and CHKV when examined by RT-PCR. Three Larval indices namely House Index, HI (percentage of houses positive for *Aedes* larvae or pupae), Container Index, CI (the percentage of containers positive for *Aedes* larvae or pupae) and Breteau Index, BI (the number of positive containers (with larvae and/or pupae per 100 inspected houses) were assessed as a measure for risk of transmission. The HI, CI and BI for both sites were as follows; Mole game reserve (HI, 42.1%, CI, 23.5% and BI, 100 for rainy season and 0 for all indices for dry season) and Larabanga (39%, 15.5% and 61 for rainy season and 2.3%, 1.3% and 2.3 for dry season). The spatial distribution of *Aedes* breeding sites in both areas indicated that *Aedes* larvae were breeding in areas with close proximity to humans. Lorry tires were the main source of *Aedes* larvae in all the study areas. Information about the species composition and the potential role of *Aedes* mosquitoes in future outbreaks of the diseases that they transmit is needed to design efficient surveillance and vector control tools.

## Introduction

Three *Aedes*-borne viral diseases, Dengue, Zika, and Chikungunya were previously known to contribute minimally to global mortality and morbidity [1]. However, over the past five (5) decades the occurrence of these diseases has increased exponentially [2]. Amongst these three diseases, Dengue infection has become the most dangerous and most rapidly spreading infection worldwide [3]. In Africa, Dengue has been reported in 34 countries including Togo, Burkina Faso, Côte d’Ivoire, Gabon and others with their capital cities being the most severely affected [4,5]. Zika virus (ZIKV) infection which is closely related to Dengue virus (DENV) infection has circulated in Africa and Asia after its discovery in Rhesus monkeys in the Zika forest of Uganda in 1947 [6]. The disease has received a striking public health attention due to its association with microcephaly and other neurological disorders such as the Guillian Barre syndrome in babies born to infected mothers [7]. The very first outbreak of Chikungunya disease occurred in Tanzania in 1953. Since then, a few outbreaks and sporadic cases were reported mainly in Asia and Africa.

The viruses that cause Dengue, Zika and Chikungunya are transmitted by mosquitoes of the *Aedes* species specifically *Aedes aegypti* and *Aedes albopictus*. All three viruses are transmitted in forest cycle (zoonotic) which involves non-human primates and arboreal mosquitoes [8]. Nonetheless, the domestication and the spread of the vectors outside their native range have caused a spillover from the zoonotic transmission cycle to urban transmission cycle involving human primates. Among the *Aedes* species, *Aedes aegypti* is known to be the most efficient vector in the transmission of these viruses [9]. They are efficient vectors because they have learnt to live their entire life cycle in close proximity to humans. They also feed solely on human blood even in the presence of other mammals as humans are the most available and stable source of blood [10].The *Aedes aegypti* species is highly susceptible to all three viruses and are efficient in propagating these diseases because during the intake of a single blood meal, they bite several humans hence transmit the virus to multiple hosts [1]. *Aedes albopictus*, also known as the “Asian tiger” mosquito, has become an important vector in many regions as it complements the role of *Aedes aegypti* in a lot of places. The spread of this invasive species of *Aedes* mosquitoes is increasing rapidly. It has become effective in transmitting some of these *Aedes*-borne viruses especially CHIKV. This is due to the E1-A226V mutation in the Eastern/Central/South African genotype which has conferred an enhanced transmission of the Virus in *Aedes albopictus* [11]. There are no vaccines for Dengue, Zika and Chikungunya infections, vector control is the most viable approach to controlling these diseases [12,13].

Ghana has previously not recorded outbreaks of Dengue, Chikungunya and Zika virus infections but the country is at risk due to its proximity to West African countries where recent outbreaks had occurred [14]. Yellow fever (YF) outbreaks had occurred in Ghana in recent years [15,16]. However, the presence of a vaccine against YF has substantially mitigated the risk and outbreaks. Recent studies have established previous exposure to Dengue virus among some diagnosed malaria patients in some urban areas in Ghana [17]. A recent study confirmed the presence of Dengue virus in the blood of two children in the Greater Accra region. Further investigations showed that these children had not travelled outside the country and therefore it can be postulated that the infection was acquired locally [18]. Local transmission may also mean that they may have acquired the infection from the bite of an infected *Aedes* mosquito. There is however very limited information on the distribution, abundance and risk of transmission of *Aedes*-borne arboviruses such as DENV, CHKV and ZIKV in the country.

The Mole Game Reserve located near Damongo in Northern Ghana has been previously described as a high-risk area for transmission of viral hemorrhagic fevers because of the presence of high population of *Aedes aegypti* in the Damongo area [19]. However, the southern part of Ghana has been found to be a low risk area [20]. The Damongo and Mole Game Reserve area have been a hotspot for Yellow fever outbreaks in recent times. The area is also a tourist site that receives many visitors from different parts of the world including some areas endemic for DENV, CHKV and ZIKV. The presence of travelers in and around the game reserve where there are primates that serve as reservoir of these viruses makes it a potential area for future outbreaks of DENV, ZIKV and CHKV. It is therefore highly imperative to conduct entomological studies in this area to better inform the relevant authorities about possible outbreaks and plan control strategies against the vectors. This study was aimed at investigating the species composition and assessing the risk of transmission of arboviral diseases around some yellow fever hotspot areas in Northern Ghana.

## Materials and Methods

### Study area and design

A cross-sectional entomological study was carried out in the Mole Game Reserve quarters area and Larabanga in the Northern region of Ghana. The survey was carried out in the rainy season (October 2018) and the dry season (March 2019). Both areas are in the West Gonja district. The Mole Game Reserve area is a guinea savannah ecological zone with latitude and longitude of 9°30’ 0” N 2°0’ 0” W with an elevation of 200 meters. The Game Reserve is home for about 400 species of animals including elephants, chimpanzees, birds and others. It attracts a great number of tourists both nationally and internationally. Larabanga (9° 13’ 0” N 1° W 51’ 0”) is 44km away from the Mole Game Reserve. The village is known for its Sahelian mosque which is the oldest mosque in all of Ghana and if possible West Africa. Tourists who visit the game reserve usually visit the Larabanga mosque due to its proximity. These sites were selected based on the potential of zoonotic infections and also previous histories of outbreaks of Yellow fever and suspected viral haemorrhagic fevers which are caused by flavivirus.

### Mosquito collection and larval survey

Adult mosquitoes were sampled using baited BG-Sentinel traps and sweep nets to capture *Aedes* mosquitoes resting on vegetation outdoors. BG-Sentinel traps were set at 06:00h and collected 12h later. Adult mosquitoes collected were grouped according to sex and put in 2ml vials and labelled with information on place and date of collection after which they were immediately stored in liquid nitrogen to keep the integrity of the genetic material. Larvae were collected from selected houses and potential breeding sites in and around the Mole Game Reserve as well as in Larabanga using ladles, dippers, pipettes and buckets. Larvae collected were grouped and stored in appropriately labelled falcon tubes containing RNA-later to keep the integrity of the genetic material and also to prevent decomposition that will change morphological characteristics. All samples were transferred to the Noguchi Memorial Institute for Medical Research-Vector laboratories where they were morphologically identified to species level using identification keys [21].

Houses and containers inside and around houses were inspected for the presence of *Aedes* larvae and pupae. The data from the larval survey was used to calculate the larval indices which are a surrogate marker for the risk of transmission of Arboviruses. During the survey sites where *Aedes* samples were collected and the houses that were inspected for immature stages of *Aedes* larvae were Geo-referenced using Geographical positioning system (GPS) and the data spatially displayed on a map.

### Molecular identification of *Aedes aegypti* species

DNA was extracted from the legs and last abdominal segment of each *Aedes* mosquito sample using the Qiagen kit following the protocol described by the manufacturer. DNA extracted was amplified using Random-amplified polymorphic DNA polymerase chain reaction (RAPD-PCR). This involved amplification of random segments of *Aedes* mosquito DNA using a 10 base pair primer B3 [22]. A volume of 20μl was used for the PCR reaction. This contained 13.2 μl *of* DNase free water, 2 μl of 10 x reaction buffer, 0.6 μl of 50uM Magnesium Chloride, 0.5 μl of 10uM dNTPs, 0.5 μl of 10uM B3 primer, 0.25 μl of uM Taq Polymerase and 3 μl of DNA template. The PCR cycling conditions includes an initial denaturation step at 94°C for 4mins, followed by 45 cycles consisting of 94°C for 1min, 35°C for 1min and 72°C for 2mins and a final extension step at 72°C for 5mins. The PCR products were electrophoresed at 100volts for 1 hour15minutes on 1.5% agarose gel stained with ethidium bromide. The PCR products were run alongside 1kb ladder and visualized using a trans-illuminator.

### Processing of mosquitoes for viral detection

Mosquitoes were grouped in pools of 30 and homogenized in Minimum Essential Medium (MEM) containing Earle’s Salts, 2% L-glutamine, supplemented with 10% heat inactivated FBS and Penicillin-Streptomycin. The homogenate were centrifuged at 12000rpm for 3min and the supernatant divided into two new tubes. One tube was labeled for storage at −80°C and the other used for RNA extraction.

### Viral detection using Trioplex Real-time RT-PCR Assay

Total RNAs of the samples were extracted using the QIAmp Viral Mini Kit according to the manufacturer’s protocol. The Trioplex Assay for detection of DENV, CHIKV and ZIKV were done using AgPath-ID RT-PCR kit following the protocol by Centres for Disease Control and Prevention [23]. The RT-PCR reaction of 20μl was made up of 0.5μl Nuclease free water, 12.5μl of 2x RT Buffer, 0.5μl of DENV mix, 0.5μl of CHIKV mix, 0.5μl of ZIKV mix, 0.5μl of enzyme mix and 5μl of RNA template. The Real-time RT-PCR cycling conditions includes a reverse transcription step at 50°C for 30mins, a hot start step at 95°C for 2mins, a denaturation step at 95°C for 15sec and an annealing and extension step at 60°C for 1mins for 45 cycles. This was done using a 7500 fast real-time machine. Nuclease free water was used as the negative control and inactivated DENV, ZIKV and CHIKV were used as positive controls.

### Data analysis

Three indices were used to assess *Aedes* mosquito density in the various collection sites. These were House index (HI), Container index (CI) and Breteau index (BI). HI was expressed as the percentage of houses infested with *Aedes* larvae; CI as the percentage of containers infected with larvae or pupae; and BI as the number of positive containers per 100 inspected houses. The risk of transmission of *Aedes*-borne viruses at the study site was estimated using the WHO criteria. For the WHO criteria, an area where BI, HI, and CI exceed 50, 35 and 20 respectively, the risk of *Aedes*-borne viruses is considered to be high; BI between 5 and 50, the density of *Ae. aegypti* is considered to be sufficient to promote an outbreak of *Aedes-*borne viral disease; an area where BI, HI and CI are less than 5, 4 and 3 respectively; it is considered to be unlikely for *Aedes*-borne virus transmission to occur. The proportions of mosquitoes by species, season and site were also calculated using Fishers exact test.

## Results

A total of 1957 *Aedes* mosquitoes were collected from the study sites of which 906 (46.3%) were adult *Aedes* mosquitoes and (1051) 53.7% were *Aedes* larvae. In the rainy season 1930 *Aedes* mosquitoes were collected and in the dry season 27 *Aedes* mosquitoes were collected (Table 1). Fishers exact test indicated that *Aedes* mosquitoes collected during the rainy season were significantly higher than *Aedes* mosquitoes collected during the dry season (P < 0.001). *Aedes aegypti* was identified in both sites during the rainy and dry seasons. *Aedes vittatus* was only identified in the Mole game reserve area during the rainy season. A total of 415 *Aedes aegypti* mosquitoes collected from both sites during the rainy season and dry season were molecularly identified to subspecies level. The two subspecies, *Aedes aegypti aegypti* and *Aedes aegypti formosus* were identified in Mole game reserve during the rainy season (Fig 2). *Aedes aegypti aegypti* was identified in Larabanga during both seasons.

**Table 1:**
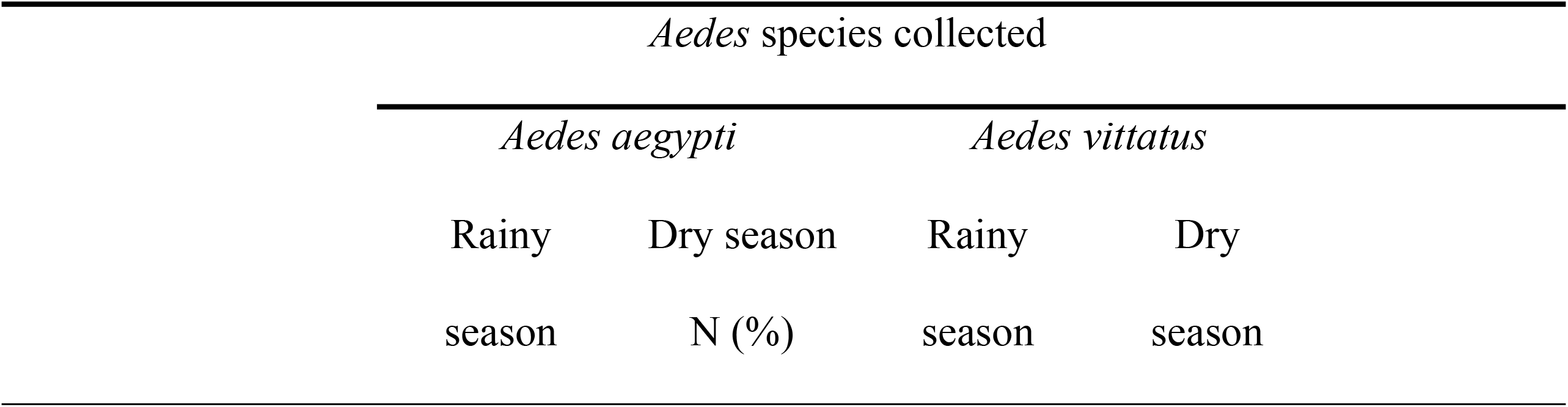
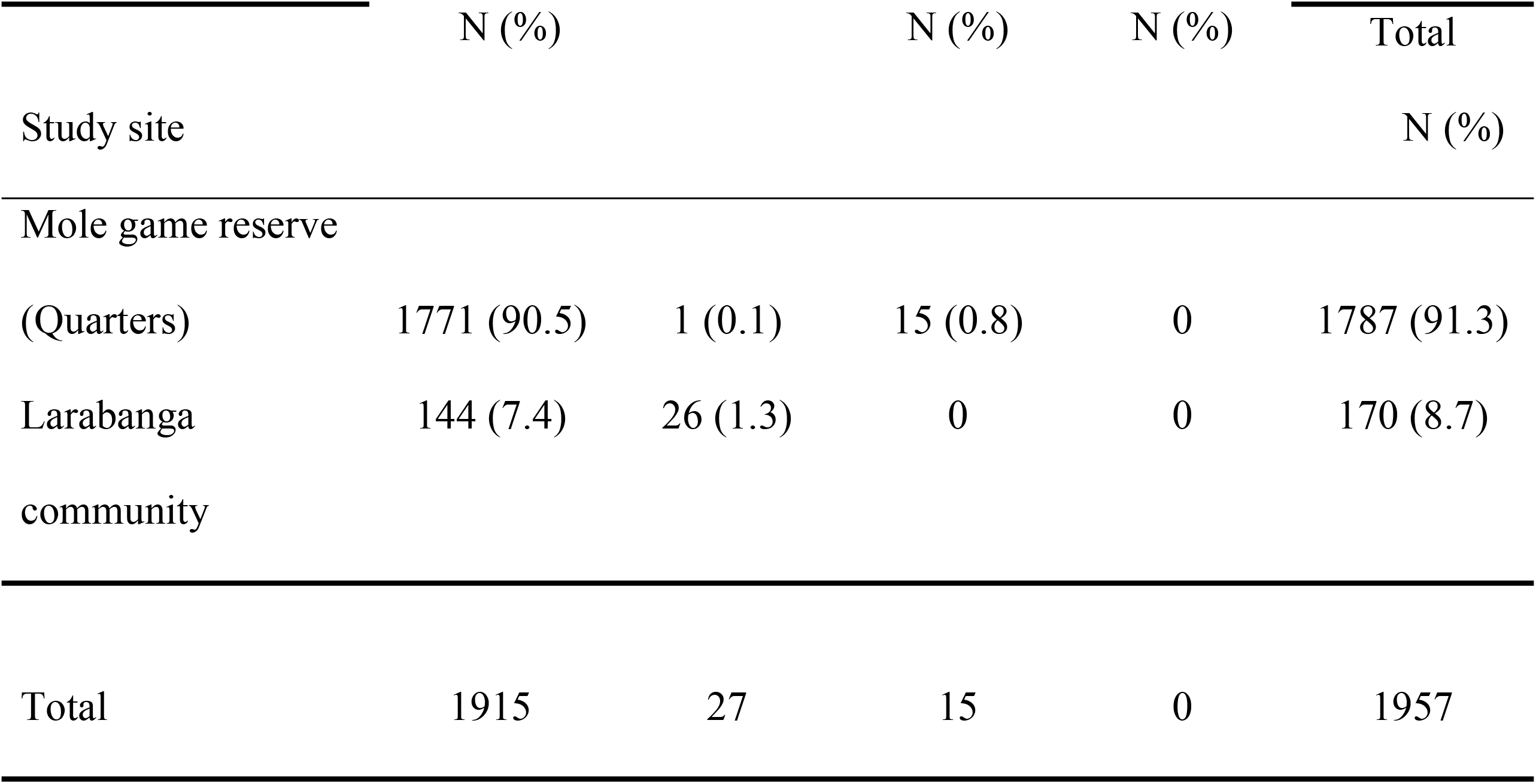
Number of *Aedes* species identified for Mole Game Reserve and Larabanga during the rainy and dry seasons.

| *Aedes* species collected |  |  |  |
| --- | --- | --- | --- |
| *Aedes aegypti* |  | *Aedes vittatus* |  |
| Rainy season | Dry season<br>N (%) | Rainy season | Dry season |

|  | N (%) |  | N (%) |  | N (%) |  | Total |
| --- | --- | --- | --- | --- | --- | --- | --- |
| Study site |  |  |  |  |  |  | N (%) |
| Mole game reserve |  |  |  |  |  |  |  |
| (Quarters) | 1771 (90.5) | 1 (0.1) | 15 (0.8) | 0 |  |  | 1787 (91.3) |
| Larabanga community | 144 (7.4) | 26 (1.3) | 0 | 0 |  |  | 170 (8.7) |
| Total | 1915 | 27 | 15 | 0 |  |  | 1957 |

**Figure 1:**
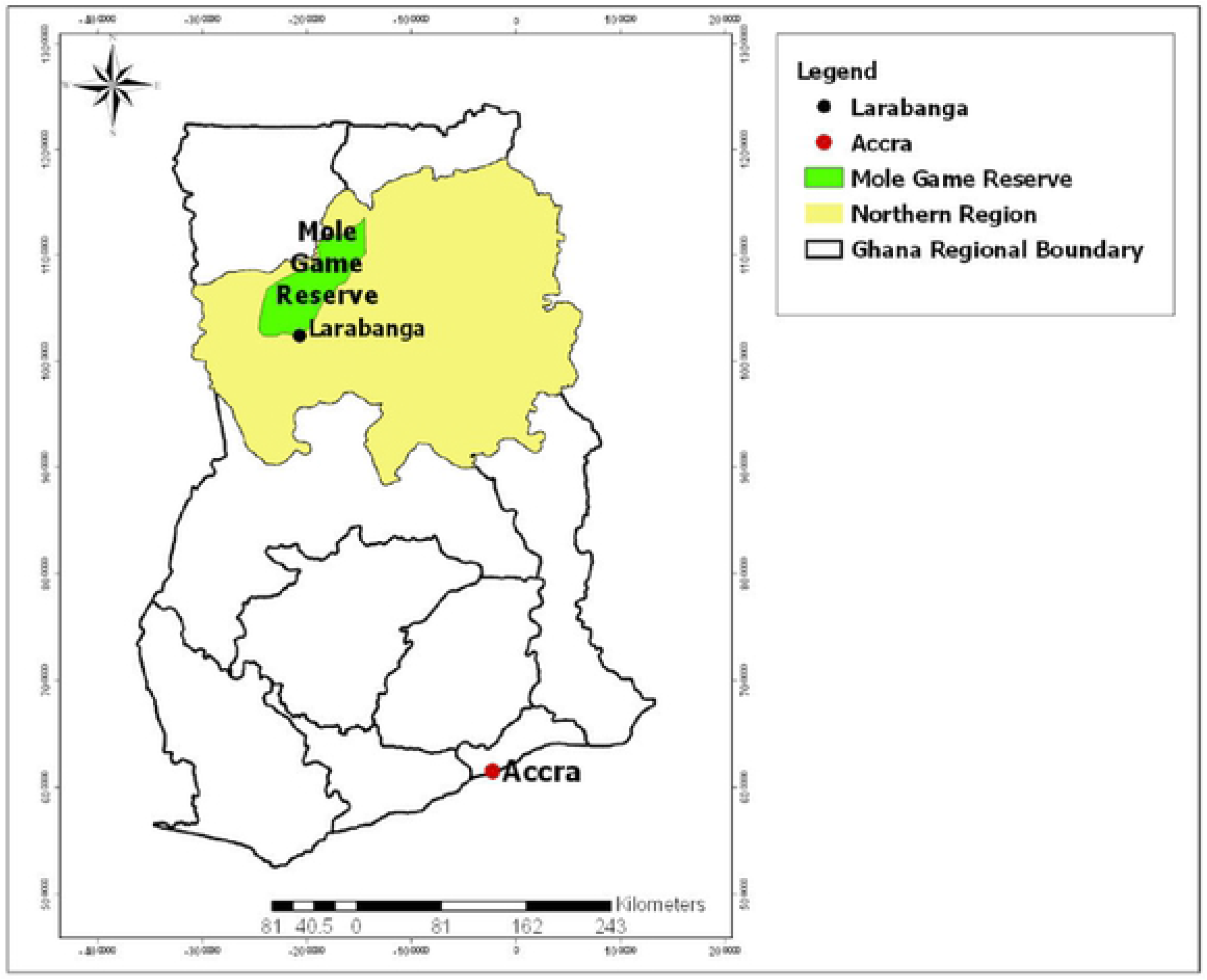
Map of Ghana showing the study sites.

**Figure 2:**
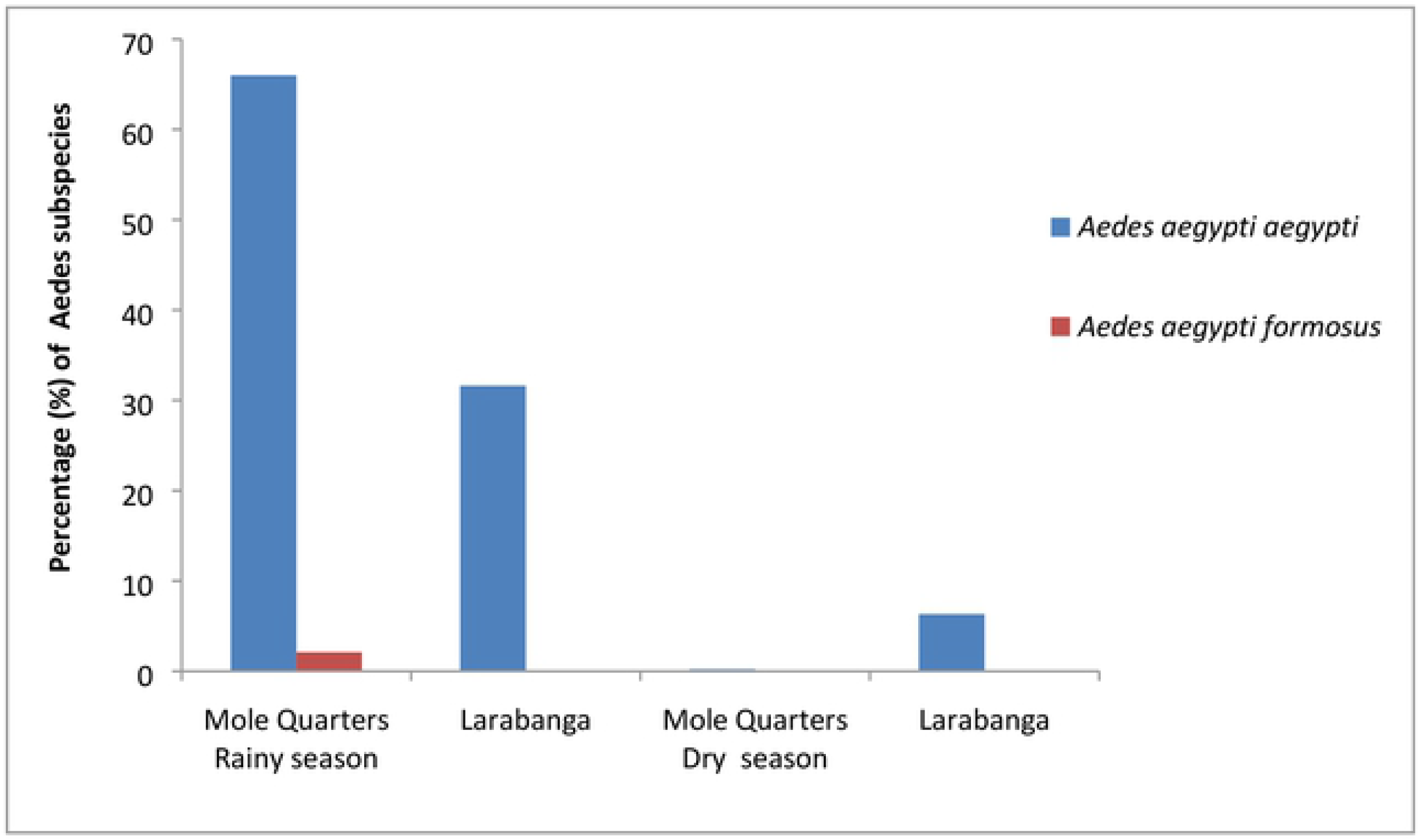
Seasonal distribution of *Aedes agypti* sub-species in the study areas

### *Aedes* larval indices

Larval indices (BI, HI and CI) were estimated for Mole game reserve and Larabanga during the rainy season and dry season. 167 households were surveyed during the rainy season and 206 households were surveyed during the dry season. All three indices were high during the rainy season than the dry season. Observed larval indices were higher in Mole quarters compared to Larabanga. All larval indices (HI, CI and BI) for Mole during the rainy season were above the WHO threshold. HI and BI were above the WHO threshold for Larabanga during the rainy season (Table 2). During the survey, 242 containers were observed of which 18.2% were positive for *Aedes* larvae during the rainy season. 220 containers were inspected during the dry season of which 2.3% were positive for *Aedes* larvae. Overall, 45.5% of households were positive for *Aedes* larvae in Mole game reserve area and 36.4% for Larabanga during the rainy season. Discarded lorry tires and ater storage containers constituted over 80% of the positive habitats surveyed. There were no positive containers observed in Mole game reserve area during the dry season.

**Table 2:** *Aedes* mosquito larval indices and WHO threshold for transmission risk of *Aedes*-borne viral diseases in Mole Game Reserve and Larabanga

| Larval indices for study areas | Rainy season | Dry season | WHO Threshold for transmission risk of VHF's |
| --- | --- | --- | --- |
| **Mole Quarters** |  |  |  |
| House index | 42.1* | 0.0 | 4-35 or above |
| Container index | 23.5* | 0.0 | 3-20 or above |
| Breteau index | 100.0* | 0.0 | 5-50 or above |
| **Larabanga** |  |  |  |
| House index | 39.0* | 2.3 | 4-35 or above |
| Container index | 15.5 | 1.3 | 3-20 or above |
| Breteau index | 61.0* | 2.3 | 5-50 or above |

\*above the WHO threshold

### Spatial distribution of *Aedes aegypti* breeding sites in relation to human habitation

In the rainy season houses positive for *Aedes* larvae were clustered just around the breeding sites. It was observed that most of the breeding sites were extremely close to households in the Mole area. There were no positive houses and breeding sites during the dry season (Fig 3&4). The same pattern was observed during the rainy season in the Larabanga area. In the dry season there were no breeding sites observed however, there were a few households that were positive for *Aedes* larvae. (Fig 5&6).

**Figure 3:**
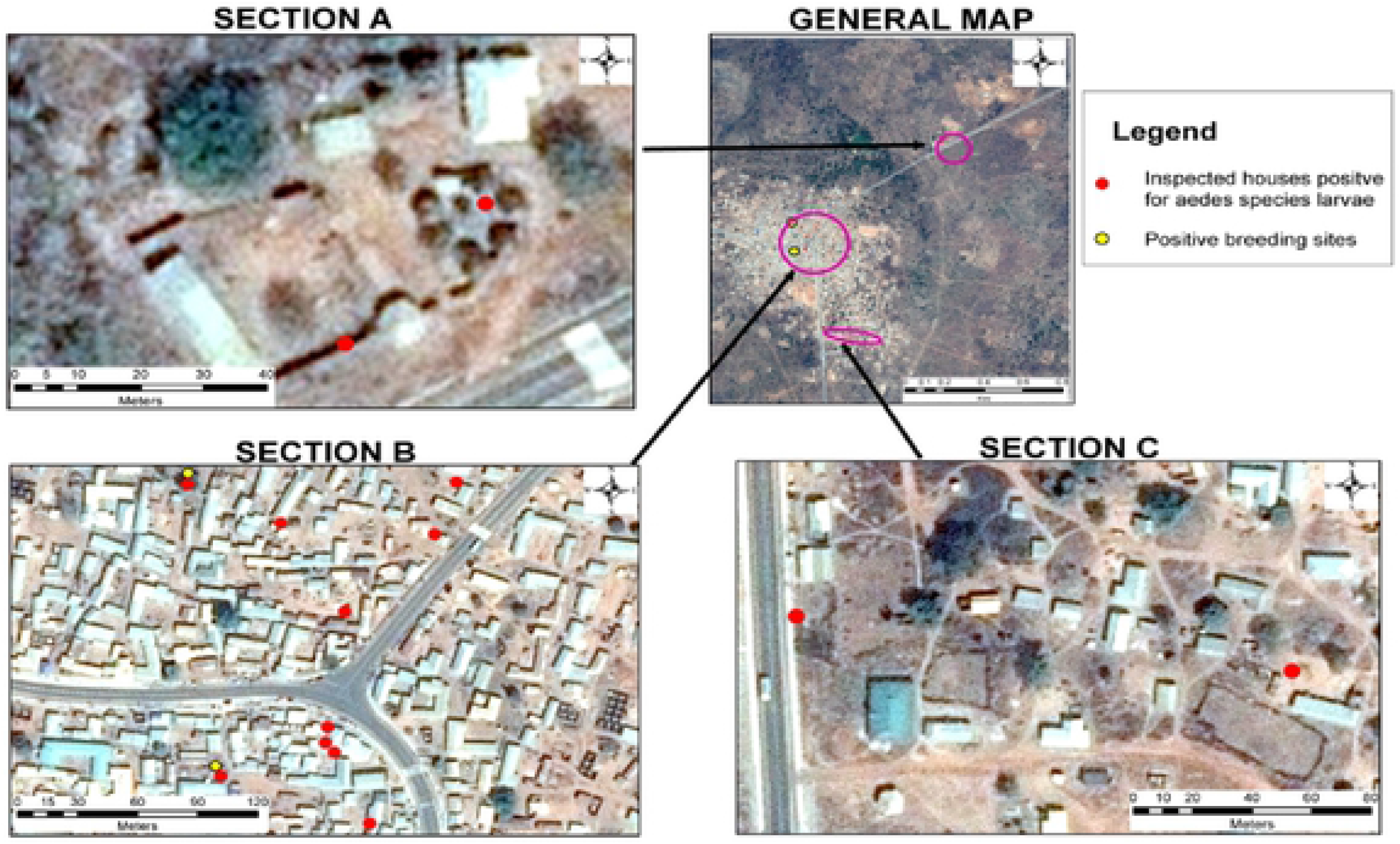
Spatial map of houses positive for *Aedes* larvae in relation to breeding sites of *Aedes* mosquitoes and human habitation in Larabanga during the rainy season.

**Figure 4:**
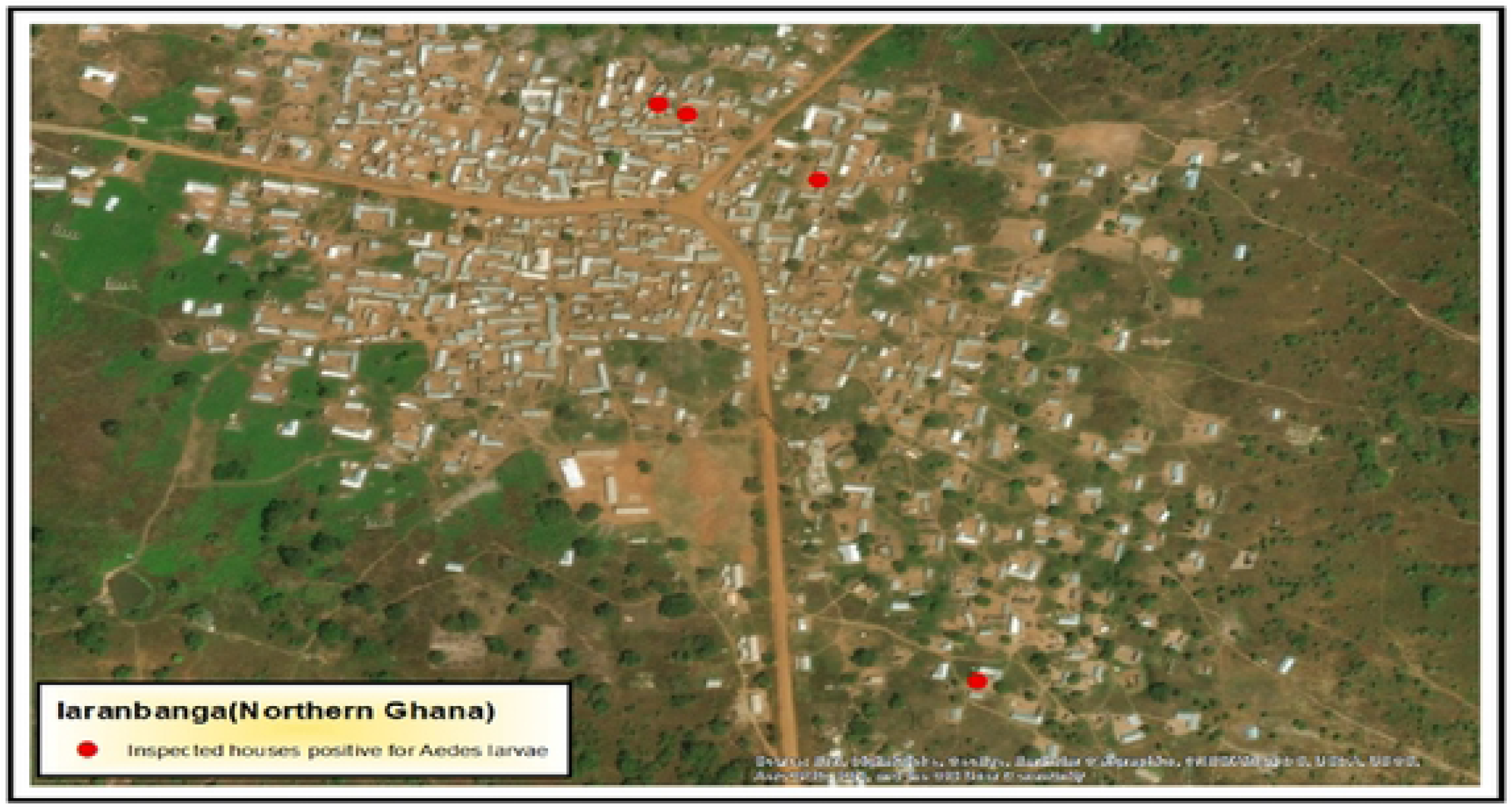
Spatial map of houses positive for *Aedes* larvae and breeding sites of *Aedes* mosquitoes and human habitation in Larabanga during the dry season.

**Figure 5:**
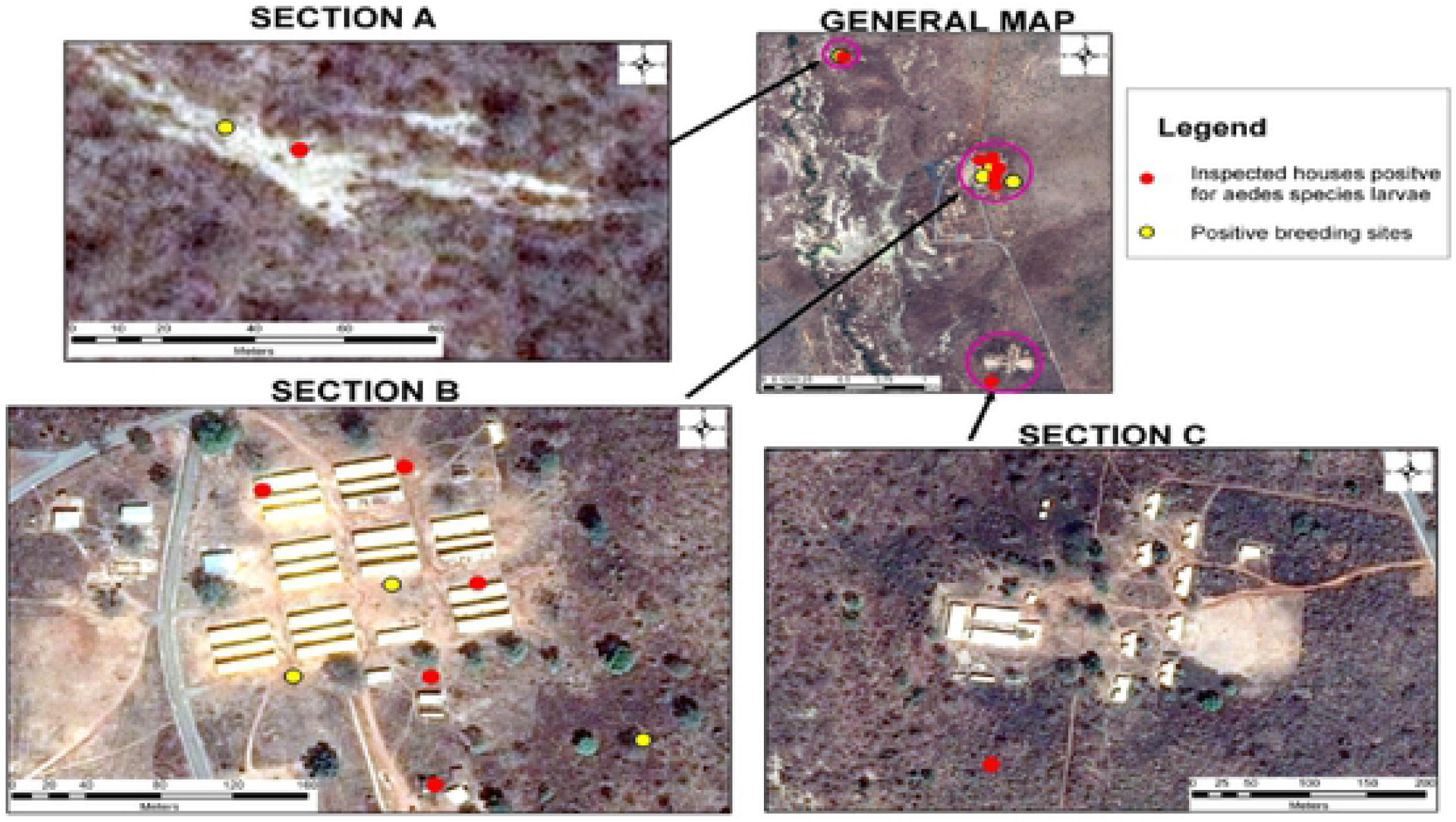
Spatial map of houses positive for *Aedes* larvae in relation to breeding sites of *Aedes* mosquitoes and human habitation in Mole Game Reserve during the rainy season.

**Figure 6:**
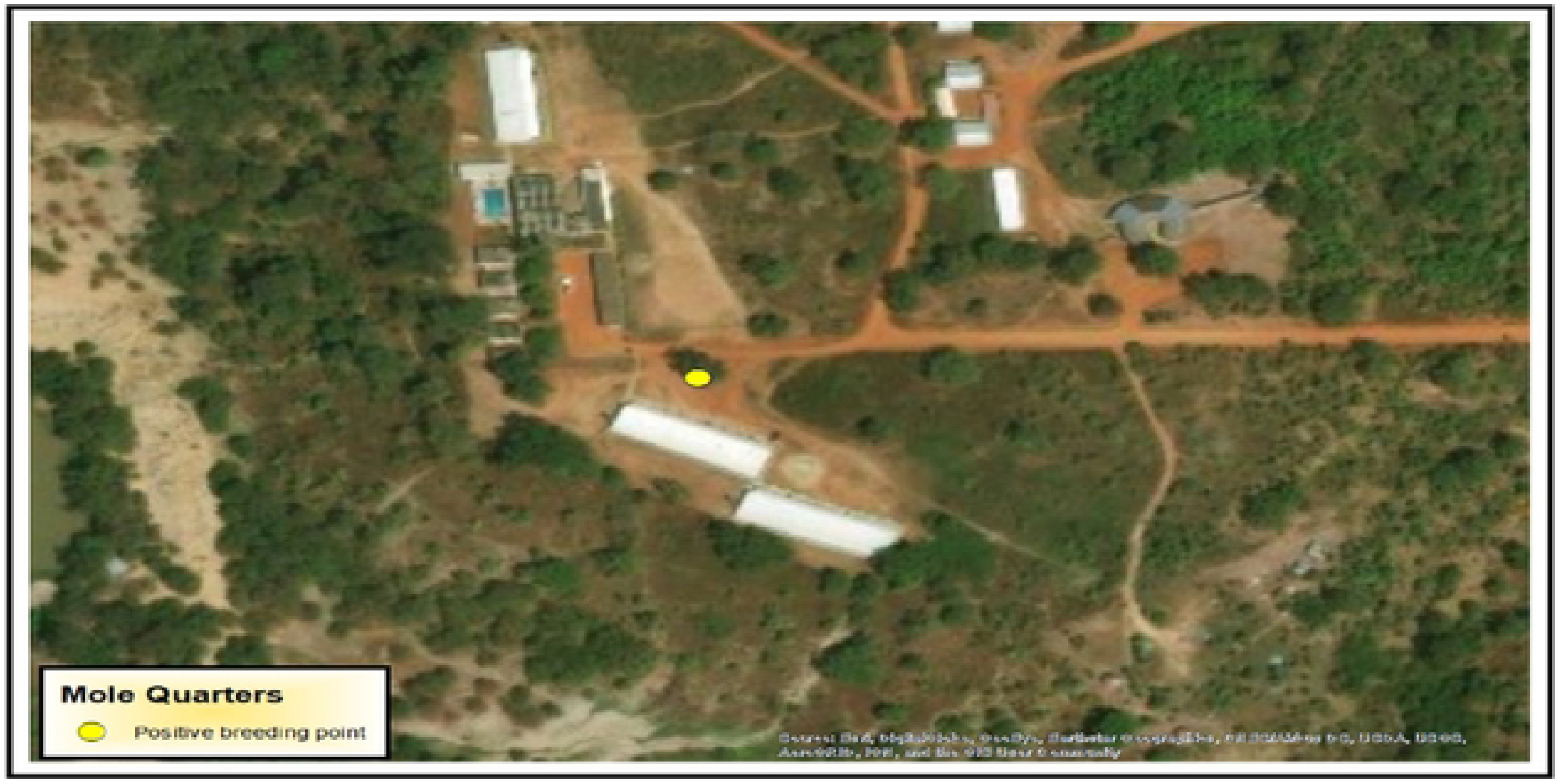
Spatial map of houses positive for *Aedes* larvae and breeding sites of *Aedes* mosquitoes and human habitation in Mole Game Reserve during the dry season

### Detections of DENV, ZIKV and CHIKV in *Aedes* samples

Overall 75 pools consisting of 1957 *Aedes* mosquitoes were tested. These included both males and females. There were 66 pools for *Aedes aegypti* and 2 pools for *Aedes vittatus* for the rainy season. *Aedes aegypti* (7 pools) collected in the dry season were also used for detection of *Aedes*-borne Arboviruses. All pools were negative for DENV, ZIKV and CHKV.

## Discussion

In this study, mosquitoes were collected from the Mole quarters and from Larabanga during the rainy and dry seasons. Larval sampling using the dipping method yielded high numbers of *Aedes* mosquitoes during the rainy season. The BG-Sentinel traps were less effective in this area during the rainy season because the traps were disrupted by rain. There were high numbers of *Aedes* mosquitoes collected during the rainy season. The abundance of *Aedes* populations is controlled by rainfall because it supports the development of additional breeding sites, hatching of eggs, growth of vegetation cover and cool shaded environment for the development of immature stages and high relative humidity [24]. *Aedes* mosquitoes collected during rainy and dry season in the study sites were predominantly *Aedes aegypti*. This may be due to the presence of discarded water holding containers and water storage containers around the human settlement which is the preferred site for *Aedes* aegypti female mosquitoes to lay their eggs [25]. This is however contrary to findings of a study conducted by Dom, Madzlan, Hasnan, & Misran (2016) where *Aedes albopictus* was the most predominant species. *Aedes aegypti* are also known to live in close proximity to humans and they prefer human blood even in the presence of other animals [10]. This is evident in the spatial maps for the sites, which shows the proximity of major breeding sites to houses positive for *Aedes* larvae. This characteristic increases the potential of transmission of arboviruses among humans. Furthermore, the high numbers of *Aedes aegypti* mosquitoes recorded during the rainy season may be due to the hatching of eggs laid by these mosquitoes during the dry season. These species of mosquitoes have their eggs being able to withstand desiccation and only hatch when conditions are favorable [27]. The presence of the major vector for the transmission of DENV, CHIKV and ZIKV in the study areas indicates the possibility of future outbreaks. The second *Aedes* species observed was *Aedes vittatus*. This species was only identified in the Mole game reserve during the rainy season. This species is usually predominant in forest and savanna areas which is a characteristic of the Game Reserve area [28]. The *Aedes vittatus* has been incriminated as a vector for yellow fever in various parts of Africa [9,28,29]. DENV, ZIKV and CHKV have been isolated from this species indicating that it has the potential to replicate and transmit these viruses experimentally [29].

The study identified *Aedes aegypti aegypti* (*Aaa*) as the dominant subspecies in both study sites during the rainy and dry seasons. This species is known to be the “domesticated” form of *Aedes aegypti* and it is closely associated with and dependent on human habitats. This may be the reason for their presence in both sites. However, the other subspecies, *Aedes aegypti formosus* (*Aaf*) was identified only in the Mole game reserve. *Aaf* are the ancestral African type of *Aedes aegypti* that prefer nonhuman mammals as a blood source [30]. The game reserve is therefore a suitable place for their survival due to the presence of a wide variety of nonhuman mammals. These species were collected from breeding sites which were close to human settlements in the game reserve area. Their presence in these areas may be due to the presence of monkeys and warthogs around human settlements.

In assessing the risk of transmission of DEN, ZIKV and CHIKV, water holding containers around the study sites were surveyed for the presence or absence of immature stages of *Aedes* mosquitoes. The presence of water holding containers around the study sites allows the breeding of *Aedes* larvae thereby increasing the population of *Aedes* mosquito and the associated risk for arbovirus transmission [31]. During the rainy season the larval indices HI, BI and CI for Mole game reserve were above the threshold values for the WHO criteria, but higher than those of a similar study in Damongo which is about 15km from the Mole Game reserve [19] where the HI, BI and CI were 87.7, 180.9 and 44.8 respectively. These high observed values for Mole Game Reserve indicates a high risk of transmission if Dengue, Chikungunya and Zika cases become established in the area.

For Larabanga, there were high values recorded for HI and BI during the rainy season. However, CI was within the threshold values. In the dry season, the larval indices for both sites were extremely low compared to the rainy season. These observations are in agreement with other studies [32,33]. However, contrasting results were observed by Appawu *et al* (2006) where it was shown that larval indices were higher in the dry season compared to the rainy season in the Northern region of Ghana. In the dry season, the Mole and Larabanga communities received pipe borne water two days in a week and most water storage containers are cleaned regularly. This may have reduced the numbers of *Aedes* mosquitoes in the area hence the low larval indices observed in those areas. The individuals in Larabanga also relied on water from the dam which was accessed daily due to the higher demand for water during the dry season. Therefore, water was not stored for longer periods during this season. In the Mole quarters area, larval indices were low during the dry season. It was observed that *Aedes* larvae were mostly found in outdoor containers than in indoor containers. This has also been observed in some studies [34] Individuals preferred storing water indoors rather than outdoors because the monkeys in the game reserve area disturbed their water collections and items when stored outdoor. Other studies showed that *Aedes* mosquitoes were breeding indoor rather than outdoor [24,35] and this may be because water containers stored outdoor are well covered, preventing *Aedes* mosquitoes from laying their eggs inside.

In this study, discarded car tires had high positivity rate for *Aedes* larvae compared to the other containers inspected during the rainy season followed by earthen ware pots. However, it was observed that the earthen ware pots were the most positive for *Aedes* larvae during the dry season. Discarded tires usually collect water and tend to harbor *Aedes* larvae without interruption. This makes it an ideal place for *Aedes aegypti* mosquitoes to breed undisturbed. Earthen ware pots are usually used to store drinking water. Due to the cool temperature, humidity and reduced light, it serves as a suitable environment for *Aedes* mosquito breeding

All pools of *Aedes* mosquitoes from Mole game reserve and Larabanga analyzed for DENV, ZIKV and CHKV were negative. This is in accordance with the report from Appawu *et al* (2006) where all *Aedes* mosquitoes collected from various sites in the Northern region of Ghana were negative for flaviviruses. This may be due to the relatively low number of *Aedes* mosquitoes collected from the area. It is recommended a longitudinal study in the area in future that may increase the sample size enough to be able to detect any virus circulating in the mosquitoes. *Aedes aegypti formosus* was one of the subspecies identified in the study area. This species has been shown to be refractory to DENV especially DENV serotype 2 [36]. This may be the reason why these viruses were not detected and possibly why there are no outbreaks of these diseases in Ghana although the area is noted for YF outbreaks. Vector competency studies is recommended in the area to determine the role of the *Aedes aegypyi aegypti* and *Aedes aegypyi formosus* subspecies in the transmission of DEN and other arboviruses in the area.

## Conclusions

In this study, *Aedes Aegypti* and *Aedes vittatus* were the two species identified around the Mole game reserve. The predominant species of *Aedes* identified in both study sites was *Aedes aegypti* which was documented in the rainy season and dry season. Among the *Aedes aegypti* species identified, majority were *Aedes aegypti aegypti*. The risk of transmission for DENV, ZIKV and CHKV in the study areas as shown by the larval indices were high during the rainy season than the dry season. This study found that discarded tires and earthen ware pots were the preferred breeding habitats for these species. The study also showed that *Aedes* mosquitoes were breeding in close proximity to human habitation and all *Aedes* mosquito pools were however negative for DENV, ZIKV and CHKV.

## Abbreviations

DENV: Dengue Virus
CHIKV: Chikungunya Virus
ZIKV: Zika Virus
CDC: Centers for Disease Control and Prevention
WHO: World Health Organization
PCR: Polymerase Chain Reaction
DNA: Deoxyribonucleic Acid
RNA: Ribonucleic Acid
RAPD-PCR: Random Amplified Polymorphic DNA Polymerase Chain Reaction
RT-PCR: Reverse Transcriptase Polymerase Chain Reaction
HI: House Index
BI: Breteau Index
CI: Container Index
GPS: Global Positioning System
NMIMR: Noguchi Memorial Institute for Medical Research
GIS: Geographical Information System

## Acknowledgements

We thank all the fieldworkers who assisted in the data collection as well as staff of Noguchi Memorial Institute for the laboratory support. The authors are grateful to Japan Agency for Medical Research and Development (AMED) for partially supporting the study as part of the collaborative project between NMIMR and Tokyo Medical and Dental University in Japan.

## Authors’ contributions

Conceptualization: Samuel Kweku Dadzie

Data curation: Joannitta Joannides, Samuel Kweku Dadzie

Formal analysis: Joannitta Joannides, Mawuli Dzodzomenyo, Samuel Kweku Dadzie

Funding acquisition: Samuel Kweku Dadzie

Investigation: Joannitta Joannides,

Methodology: Samuel Kweku Dadzie, Joannitta Joannides

Project administration: Samuel Kweku Dadzie

Resources: Samuel Kweku Dadzie

Supervision: Samuel Kweku Dadzie, Mawuli Dzodzomenyo

Validation: Samuel Kweku Dadzie

Writing – original draft: Joannitta Joannides

Writing – review & editing: Mawuli Dzodzomenyo, Maxwell A. Appawu, Kofi Bonney, Rebecca Pwalia, Samuel Kweku Dadzie

JJ designed the study, participated in the field sample collection, performed laboratory and data analyses, and prepared the manuscript for publication. SKD designed the study, supervised the study and reviewed the manuscript. MD, MAA, KB, RP supervised and reviewed the manuscript. DB, KK, RR, AB, HT reviewed the manuscript. JON, FA, GKA, DP and EEA assisted in Laboratory experiments and sample collection.

